# A time-lagged association between the gut microbiome, nestling weight and nestling survival in wild great tits

**DOI:** 10.1101/2020.09.30.320804

**Authors:** Gabrielle L. Davidson, Shane E. Somers, Niamh Wiley, Crystal N. Johnson, Michael S. Reichert, R. Paul Ross, Catherine Stanton, John L. Quinn

## Abstract

1. Natal body mass is a key predictor of viability and fitness in many animals. While variation in body mass and therefore juvenile viability may be explained by genetic and environmental factors, emerging evidence points to the gut microbiota as an important factor influencing host health. The gut microbiota is known to change during development, but it remains unclear whether the microbiome predicts fitness, and if it does, at which developmental stage it affects fitness traits.
2. We collected data on two traits associated with fitness in wild nestling great tits (*Parus major*): weight and survival to fledging. We characterised the gut microbiome using 16S rRNA sequencing from nestling faeces and investigated temporal associations between the gut microbiome and fitness traits across development at day 8 (D8) and day 15 (D15) post-hatching. We also explored whether particular microbial taxa were ‘indicator species’ that reflected whether nestlings survived or not.
3. There was no link between mass and microbial diversity on D8 or D15. However, we detected a time-lagged relationship where weight at D15 was negatively associated with the microbial diversity at D8, controlling for weight at D8, therefore reflecting <u>relative</u> weight gain over the intervening period.
4. Indicator species analysis revealed that specificity values were high and fidelity values were low, suggesting that indicator taxa were primarily detected within either the survived or not survived groups, but not always detected in birds that either survived or died. Therefore these indicator taxa may be sufficient, but not necessary for determining either survival or mortality, perhaps owing to functional overlap in microbiota.
5. We highlight that measuring microbiome-fitness relationships at just one time point may be misleading, especially early in life. Instead, microbial-host fitness effects may be best investigated longitudinally to detect critical development windows for key microbiota and host traits associated with neonatal weight. Our findings should inform future hypothesis testing to pinpoint which features of the gut microbial community impact on host fitness, and when during development this occurs. Such confirmatory research will shed light on population level processes and could have the potential to support conservation.

## Introduction

Understanding the numerous factors that drive fitness has long been the focus of research in ecology and evolution (Merila, 1996; Pickett et al., 2013). This has primarily included behavioural factors, such as foraging (Patrick & Weimerskirch, 2014) and parental care (Schwagmeyer & Mock, 2008), and ecological factors, for example, food availability (Van Noordwijk et al., 1995), predation (Götmark, 2002), and more recently climate change (Blomberg et al., 2014). Proximate factors such as genetics (e.g. Merila (1996), Norris (1993)), stress (Crino & Breuner, 2015; Weber et al., 2018) and natal environment effects (Keller & Van Noordwijk, 1994; Pickett et al., 2013) have also been studied intensively. More recently, a growing body of research is beginning to show the gut microbiome – the microbial community in the gut and the host environment - as a powerful proximate mechanism influencing host health and fitness among animals (Kinross et al., 2011; Lloyd-Price et al., 2016; Rosshart et al., 2017). Emerging evidence has highlighted that the gut microbiota varies substantially within and between individuals (David et al., 2014; Maurice et al., 2015), and that microbe-host interactions are likely to be taxonomically widespread and central to many behavioural, ecological and evolutionary processes (Alberdi et al., 2016; Davidson et al., 2020; Sherwin et al., 2019). However, the role of the gut microbiome in determining fitness in natural populations, and whether there are critical developmental windows whereby it affects future weight and survival, remains poorly understood.

Evidence suggests that the microbiome is associated with many aspects of survival and fitness. In humans, the maternal vaginal microbiota is associated with pre-term birth (Fettweis et al., 2019) and sudden infant death syndrome (Fettweis et al., 2019; Highet et al., 2014), and the gut microbiome is associated with longevity (Biagi et al., 2016). Laboratory-bred mice, which have substantially different gut microbiota than their wild counterparts, were more likely to survive an influenza infection if they had received a gut microbiota transplant from wild donors (Rosshart et al., 2017). Notably, wild mice had higher abundances of Bacteroidetes and Proteobacteria, and lower abundances of Firmicutes, suggesting these taxa may play a role in regulating host health. The potential for the gut microbiome to impact animal fitness in natural systems has gained much attention in recent years (Amato, 2013; Suzuki, 2017; Trevelline et al., 2019; West et al., 2019). However, the empirical evidence to support such hypotheses is lacking, perhaps owing to the logistical constraints associated with collecting longitudinal data on both the gut microbiome and indicators of fitness such as survival in these systems.

Research on laboratory model species has shown that the microbial community in the gut can confer specific health benefits to the host (Kinross et al., 2011), and the timing of gut colonization is critical for normal development (Cox et al., 2014; Hansen et al., 2012). In particular, gut microbiota are important for the development and regulation of gut immunity by maintaining intestinal barrier function (Hooper et al., 2012; O’Hara & Shanahan, 2006) and aiding digestive capabilities through the metabolism of dietary components (Rowland et al., 2018). Gut bacteria can induce expression of host genes that, for example, code for proteins in the mucosal layer of the intestines (O’Hara & Shanahan, 2006). Moreover, bacterial genomes can produce enzymes that are absent in the host, yet are necessary for specific digestive functions such as cellulose digestion in termites (*Nasutitermes* spp.) and giant pandas (*Ailuropoda melanoleuca*) (Warnecke et al., 2007; Zhu et al., 2011) and nitrogen metabolism in birds (Vispo & Karasov, 1997). Germ-free mice devoid of any microbes have depressed immune function and lower nutrient assimilation efficiency, which can only be improved by introducing intestinal microbes from normal mice donors (Hansen et al., 2012; Olszak et al., 2012). In many cases, the timing of microbial colonization is critical and microbiome restoration must occur within an early developmental window in order to affect host phenotypes that are observed later in life (Cox et al., 2014; Hansen et al., 2012). Therefore, variation in host microbiota can have important consequences for digestive capability and body condition, and act as a mechanism underlying developmental plasticity (Alberdi et al., 2016). Importantly, effects may be dependent on the timing of microbial colonization if microbial effects on host traits may have immediate effects (i.e. contemporary associations), or delayed effects (i.e. time-lagged associations).

In many vertebrate taxa, natal body size (Bowen et al., 2015; Ringsby et al., 1998; Stige et al., 2019), natal body mass (Both et al., 1999; Monrós et al., 2002), or growth rate (Martin, 2015; Mikaela et al., 2002) can be key predictors of juvenile viability and recruitment into the breeding population. The association between the gut microbiome and weight has been studied extensively in clinical studies for application in human obesity (Bäckhed et al., 2004; Turnbaugh et al., 2006), and in agricultural studies to increase animal productivity (Coates et al., 1981; Ford & Coates, 1971; Torok et al., 2011). While having a more diverse microbiome may intuitively suggest a healthier gut microbiome (Lozupone et al., 2012), the effects of microbial community structure are likely more complex (Zaneveld et al., 2017). In humans, lower diversity of gut microbiota is associated with high body mass index and high amounts of visceral fat (Beaumont et al., 2016; Le Chatelier et al., 2013). Specific taxa, such as *Lactobacillus* spp. can influence fat storage and weight change, however, the effects of such bacteria are strain-specific (Drissi et al., 2017). In poultry, beneficial bacteria can be selectively enriched (e.g. Patterson et al. (1997)), and the use of antibiotics reduce gut bacterial load, decrease microbial diversity, and improve feed conversion ratios (Banerjee et al., 2018). Weight gain associated with antibiotic administration and decreased bacterial diversity has also been shown in house sparrows (*Passer domesticus*) (Kohl et al., 2018), Daphnia magna (Motiei et al., 2020) and magellanic penguins (*Spheniscus magellanicus*) (Potti et al., 2002). By contrast, microbial diversity has been positively associated with body condition and growth rate in wild nestling birds (Teyssier, Lens, et al., 2018).

The great tit (*Parus major*) provides a powerful natural system for addressing microbiome related effects on host phenotypes. It is an altricial species that readily breeds in nest boxes, so that the developmental period can be tracked easily in the wild. We investigated the relationship between natural variation in the gut microbiome and direct and indirect measures of fitness-related traits in nestling great tits when they were 8 and 15 days old. The neonatal stage is a critical time for gut microbiome colonization (Koenig et al., 2010) and the microbial-programming of adult phenotypes (Cox et al., 2014; Hansen et al., 2012; Sudo et al., 2004). Nestling great tits undergo substantial shifts in gut microbiota between day 8 and day 15 of development (Teyssier, Lens, et al., 2018), a period when we predicted the microbiome would have a detectable effect on body weight and the probability of survival to fledging. We tested whether diversity of the microbiome, or the abundances of specific microbes, were associated with nestling weight and survival to fledging. Given the mixed reports of microbial links with host fitness in the existing literature (see above), we had no a priori predictions regarding the direction of such potential relationships. Nevertheless, we proposed that if diversity was positively related to measures of host fitness, then a rich and/or evenly distributed microbial community would be important for healthy development. By contrast, if the relationship was negative, then the host may benefit from the presence or absence of specific microbial taxa, which should be observed in differences in the relative abundance of these specific microbes associated with host fitness. Moreover, because early developmental windows have been reported to be important for microbial influences on host traits (Cox et al., 2014; Hansen et al., 2012), we sampled the gut microbiome from chicks at two developmental stages to test whether microbiome effects on host fitness traits were contemporary or time-lagged. In other words, we asked whether the gut microbiome at D8 or D15 was a better predictor of mass and survival.

## Methods

### Field monitoring and microbiome sampling

Birds were sampled from nine nest box populations across Co. Cork, Ireland, five of these were mixed/deciduous habitats and four were coniferous habitats (see O’Shea et al. (2018)). We collected 204 faecal samples from 150 nestling great tits from 54 nests (see below for the number of samples that were successfully sequenced and sample size per developmental stage) for 16S rRNA gene sequencing in order to map the gastrointestinal microbial communities. During April-June 2016, nest boxes were monitored to determine lay dates, hatching dates and nestling survival. Nestlings aged 8 days (+/− 1 day) (D8) and 15 days (D15) were placed into sterile holding bags inside a heated holding case. Coffee filters were used to line the bags in order to soak up uric acid from the faeces, as uric acid has the potential to affect downstream sequencing (Khan et al., 1991). Samples were collected from nestlings that defaecated naturally within 15-20 minutes of being out of the nest before being returned to the nest. Although the aim was to have repeated samples for all individuals, not all birds survived, and not all birds produced samples within this time limit. Faecal sacks were opened using a sterile inoculation loop to release the faecal matter and place in a microcentrifuge tube containing 500uL of 100% ethanol. Samples were stored at −20°C within 8 hours of collection until DNA extraction. D8 birds were weighed, and uniquely individually identified by clipping the tip of one of their nails, avoiding the blood vessel (commonly known as the “quick”). D15 nestlings were weighed and ringed with a unique identifiable metal ring (British Trust for Ornithology) and matched against their unique nail clipping. Nestlings for which we had repeated samples (i.e. for both D8 and D15) are referred to as ‘repeat samples’.

### DNA Extractions

Briefly, DNA was extracted from the dried faecal contents of all birds using the Qiagen QIAamp DNA Stool Kit, following the “Isolation of DNA from Stool for Pathogen Detection” protocol (June 2012 edition), with modifications described in (Shutt et al., 2020) to accommodate dried avian faeces. A 0.10 - 0.20 g aliquot of each faecal sample was added to the kit, alongside two negative controls which were carried through to sequencing. The negative controls had low sequence reads and were not included in any subsequent analyses. We note that no decontamination methods were used, though they can be beneficial for samples with low biomass (Eisenhofer et al., 2019). Therefore, we recognise that our sequence data may contain some Amplicon Sequence Variants (ASVs) from the external environment, rather than the gut microbiome, although this should not have systematically biased our results. DNA was stored at −20°C. Full extraction methods are described in Supporting Information.

### Illumina MiSeq sequencing

Full library preparation details are described in Supporting Information and in Davidson et al 2020b). Briefly, the V3-V4 variable region of the 16S rRNA gene was amplified from the DNA extracts using the 16S metagenomic sequencing library protocol (Illumina). The DNA was amplified with primers specific to the V3-V4 region of the 16S rRNA gene which also incorporates the Illumina overhang adaptor. Samples were sequenced on the MiSeq sequencing platform (Clinical Microbiomics, Denmark), using a 2 × 300 cycle kit, following standard Illumina sequencing protocols.

### Bioinformatics and statistical analysis

The DADA2 pipeline (Callahan et al., 2016) was used to process the raw sequencing data in R version 3.5 (R Core Team, 2019). Sequence quality was visually inspected. Sequences were trimmed to remove adapters and lower quality reads (median quality scores below 25-30 threshold) at the extremities of the sequence and filtered to remove sequences with higher than expected errors. Read errors were estimated before dereplication. Forward and reverse reads were merged to construct ‘contig’ sequences, these were used to construct a sequence table of ASVs, which counts the number of times each unique sequence is detected. The previous steps were performed for each run separately. Then the separate sequence tables were merged and chimeras removed using the ‘consensus’ method. Taxonomy was assigned to each ASV by RDP’s Naive Bayes Classifier (Wang et al., 2007) against the Silva reference database (version 132) (Quast et al., 2012). This is at 100% sequence identity in contrast to the lower resolution OTU method which groups sequences at 97% identity. ASVs allow greater sensitivity and specificity, better discrimination of ecological patterns than OTU’s and are reusable across studies (Callahan et al., 2017).

The DADA2 outputs were assembled into a single Phyloseq object (McMurdie & Holmes, 2013). Sequences identified as mitochondrial or chloroplast were removed. Further filtering of samples and ASVs took place in R. Sample completeness curves were plotted using vegan (Oksanen et al., 2019) and helped determine the lower cut-off for sample reads at 10,000 reads. Low read samples (<10000 reads, 9 samples) were removed leaving 195 (Day-8=81, Day-15=114, repeat samples=41) samples for the analysis. Alpha diversity (both Shannon and Chao1 diversity) was calculated using the ‘*estimate_richness*’ function from the phyloseq package on the filtered dataset. Shannon diversity (Shannon, 1948) measured richness weighted by abundance (the evenness of a community) and Chao1 (Chao, 1984) measured richness, specifically estimating taxa abundance and rare taxa missed from under sampling.

## Statistical analysis

### Nestling weight: Alpha diversity and phylum-level relative abundance

Linear mixed models (LMMs) and generalised linear mixed models (GLMMs) were used to examine the association between measures of nestling weight and two measures of alpha diversity: Shannon’s index and Chao1 index. Models were fit using the lme4 package (Bates et al., 2015) on each data subset. Numeric variables were centered and scaled. Model diagnostics were assessed using the DHARMa package (Hartig, 2019). Raw weight (g) was used as the response variable instead of change in weight (delta-weight) as this allowed us to control for D8 weight in time-lagged models, as well as to differentiate between, for example, very light D8 nestlings achieving normal D15 weight and heavy D8 nestlings achieving heavy D15 weight. We also tested whether alpha diversity at D8 was correlated with alpha diversity at D15 (for repeated samples within individuals) by calculating Pearson’s correlation coefficient with the ‘cor.test’ function in R.

We examined whether nestling body weight was influenced by the gut microbiome in three different ways: (1) whether the microbiome at an early developmental stage predicted future host weight (henceforth ‘time-lagged weight’), while controlling for weight at D8, (2) whether the microbiome at the time of sampling was associated with weight at the same developmental stage (henceforth ‘contemporary weight’) at D8, and (3) contemporary weight at D15. Due to collinearity (alpha diversity) and non-independence (Phylum-level relative abundance), we performed separate analyses for four different measures of microbiome as explanatory variables: alpha diversity (Shannon diversity and Chao1 index), and phylum-level abundance (Proteobacteria and Firmicutes). These two phyla were selected for our analyses because they were the only two taxa that were prevalent across all birds (except for three birds).

For the time-lagged weight analyses, we ran a linear mixed model (LMM) using the lmer function in R with D15 weight as the response variable, and alpha diversity at D8 as a fixed variable. We included D8 weight as a covariable to control for variation in starting weight. We also included habitat, lay date of first egg and brood size as fixed factors as these may influence nestling weight. Site and nest ID were used as nested random effects, as well as sequence plate as a separate random effect. Only birds that had repeat samples were included in this analysis (n= 41).

For the D8 contemporary weight analyses (all D8 samples, n= 81), we ran a LMM as described above, with D8 as the response variable and alpha diversity at D8 as a fixed factor. We included the same fixed effects and random effects as described above. The D15 contemporary analyses (all D15 samples, n = 114) were the same as described for D8, using D15 data as opposed to D8 data. D15 weight was squared in the contemporary D15 model as the model did not converge with raw weight as the response.

### Nestling survival: Alpha diversity and phylum-level relative abundance

We tested whether microbiome diversity at D8 predicted nestling survival. Here we define survival as surviving to fledgling. Birds that were previously alive at D8, but were either dead in the nest, or absent (e.g. eviction from the parents) at D15 were scored as not surviving. Nests were checked again approximately 25 days after hatching, when birds were expected to have fledged. Birds found dead in the nest were scored as not surviving, and those that were absent were assumed to have fledged because chicks were too large to be evicted, and we did not see any evidence of predation (woodpeckers are absent in our woodlands, and predation from stoats always led to the entire nest failing, and was isolated to a single woodland site not included in the current study). We ran a non-parametric cox proportional hazards survival model using the Coxme package (Therneau, 2020), and included weight (at D8), lay date and brood size (at D8) as covariables. We included three ranks of survival times (i.e. died before D15, died after D15 or fledged) as the ranks of survival times provide additional temporal information beyond a binary outcome, i.e. died/survived for the model to estimate mortality risk. The habitat variable was excluded from the survival models as it caused premature model convergence due to a lack of variation in the variable (i.e. all coniferous birds survived). Because all but one bird that survived to D15 survived to fledging, we only performed this analysis for birds that were sampled for gut microbiome at D8. We repeated this analysis with Proteobacteria abundance as the predictor variable, and again with Firmicutes abundances as the predictor variables.

### Nestling survival: Beta diversity

We investigated whether microbial community structure differed between birds that did or did not survive to fledgling, using the adonis function (PERMANOVA) from the vegan package (Oksanen et al., 2019). Beta diversity at D8 was measured using Bray-Curtis distance (weighted by ASV abundance) and Jaccard (presence/absence of ASVs). We ran separate analyses for each beta diversity measure in which D8 weight and survival (Yes/No) were fixed factors. Adonis fits each term sequentially, therefore we used the model formula *beta-diversity~D8-weight + survival*, allowing us to estimate the variation in microbiome community composition associated with survival status after accounting for body weight, as weight was associated with survival in our population (see results, Table 2). Permutations were constrained to within nest with the ‘strata’ option to control for pseudo-replication of samples from the same nest. Adonis results were very similar with and without this strata option.

The beta diversity dataset (D8 samples) was filtered by prevalence prior to distance calculation, where taxa with less than 1 copy in 5% of samples were removed following Knowles et al. (2019). This resulted in a dataset containing 1254 ASVs. The relative abundance of each phylum was calculated per sample using ‘total sum scaling’ (TSS) by dividing each feature count by the total library size (McKnight et al., 2019). Bray Curtis and Jaccard distances were calculated from our TSS normalized data. An assumption of the adonis test is that groups have homogeneity of variance. The dispersion of the groups was checked for homogeneity of variance using the Betadisper and permutest functions from the vegan package.

The beta diversity (Bray-Curtis) of D8 nestlings was visualised using a Principle Coordinate Analysis (PCoA) plot, created using the plot_ordination command in the phyloseq package. This plot compared individuals that survived to those that did not survive (see Figure S2).

### Indicator ASVs for nestling survival

We carried out indicator species analysis to explore whether ASVs were representative of birds that were alive at D8 (n=81) and survived to fledgling (n=64), or did not survive to fledgling (n=17). This analysis used the D8 data subset, which was not filtered by prevalence and contained 29709 ASVs. Indicator analysis identified “indicator species” if they reflected the state of the environment (i.e. likelihood the bird will survive, or not survive). We used the multipatt function from the IndicSpecies package (De Caceres & Legendre, 2009), which assigns a test statistic, or “indicator value” to each ASV for each group, in our case two groups: Survival-Yes, Survival-No. Taxa with an indicator value of > 0.4 and p <= 0.05 were considered indicator species. We also report the “specificity value” and the “fidelity value”. The former is the estimate of the probability that an ASV is detected in a particular group. For example, a value of 1.0 would indicate that the ASV is only detected in that group and not another group. The latter is the probability estimate of finding the species in sites (i.e. birds) belonging to the group. For example, a value of 1.0 would indicate that the ASV is recorded in all birds within that group (De Caceres, 2020). We did not adjust for multiple comparisons for three reasons. First, our analysis did not include any overlap, and thus multiple analyses, across different combinations of groups (De Caceres & Legendre, 2009). Second, the purpose of this analysis was exploratory in order to generate, rather than confirm hypotheses (Bender & Lange, 2001; Ranstam, 2019). As such, we report ASVs as potential indicator species, as opposed to reporting a quantitative, total number of taxa associated with survival/non-survival (De Cáceres et al., 2010). Third, this analysis served to generate a posteriori predictions regarding the relationship between indicator taxa and weight gain (see below), although we note that independent studies will be necessary to confirm these exploratory analyses (Ranstam, 2019). We generated a heatmap of indicator species for survival and non-survival using the heatmap function from the phyloseq package (McMurdie & Holmes, 2013).

### A posteriori analysis of indicator species

Indicator species from our analysis may reflect whether a bird survives or not. We hypothesized that indicator species may reflect host survival because such indicator species may also predict time-lagged, relative weight gain. We performed a posteriori LMMs with D15 weight as the response variable and D8 relative abundance (of the indicator species) as a fixed variable. D8 weight, habitat, brood size and lay date were also included as fixed variables. Site and nest were included as nested random terms (i.e. 1|site/nest). We restricted this a posteriori analysis to one indicator species that had the highest test statistic from each group (Survival-Yes: Lactobacillus (ASV46) and Survival-No: Rahnella (ASV38)). Because there were a high number of zeros in relative abundance of these taxa, and because our data were proportions and therefore could not be fit using a zero-inflated (count data) model, we included only birds where the taxa were present and the birds with 0 abundance for that taxa were excluded (ASV46: n=21/41; ASV38: n=8/41).

All graphs were created using ggplot2 (Wickham, 2016), unless otherwise indicated.

## Results

### Sequencing results

There were 18,537,795 high quality reads, clustered into 49,655 ASVs before removing control samples, mitochondria and chloroplast sequences. Mean reads per sample was 88,698 and individual samples ranged from 61 to 579,864 reads. Samples with less than 10,000 reads were discarded before further analysis (n=9). There were 15,183,675 total high-quality reads, clustered into 47,400 ASVs after removing control samples as well as mitochondria and chloroplast sequences. Samples ranged from 10,220 to 557,336 reads and mean reads per sample was 77,865.

The most prominent phyla across all samples (D8 and D15) were as follows (mean percentage relative abundance ±SE): Firmicutes (43.0%±2.5); Proteobacteria (35.2%±2.2); Tenericutes (9.7%±1.8); Bacteroidetes (4.2%±0.4) and Actinobacteria (2.2%±0.2). Relative abundance of the major phyla at D8 (at least >=10% relative abundance in at least 1 sample) per sample are plotted in Figure 1.

**Figure 1.**
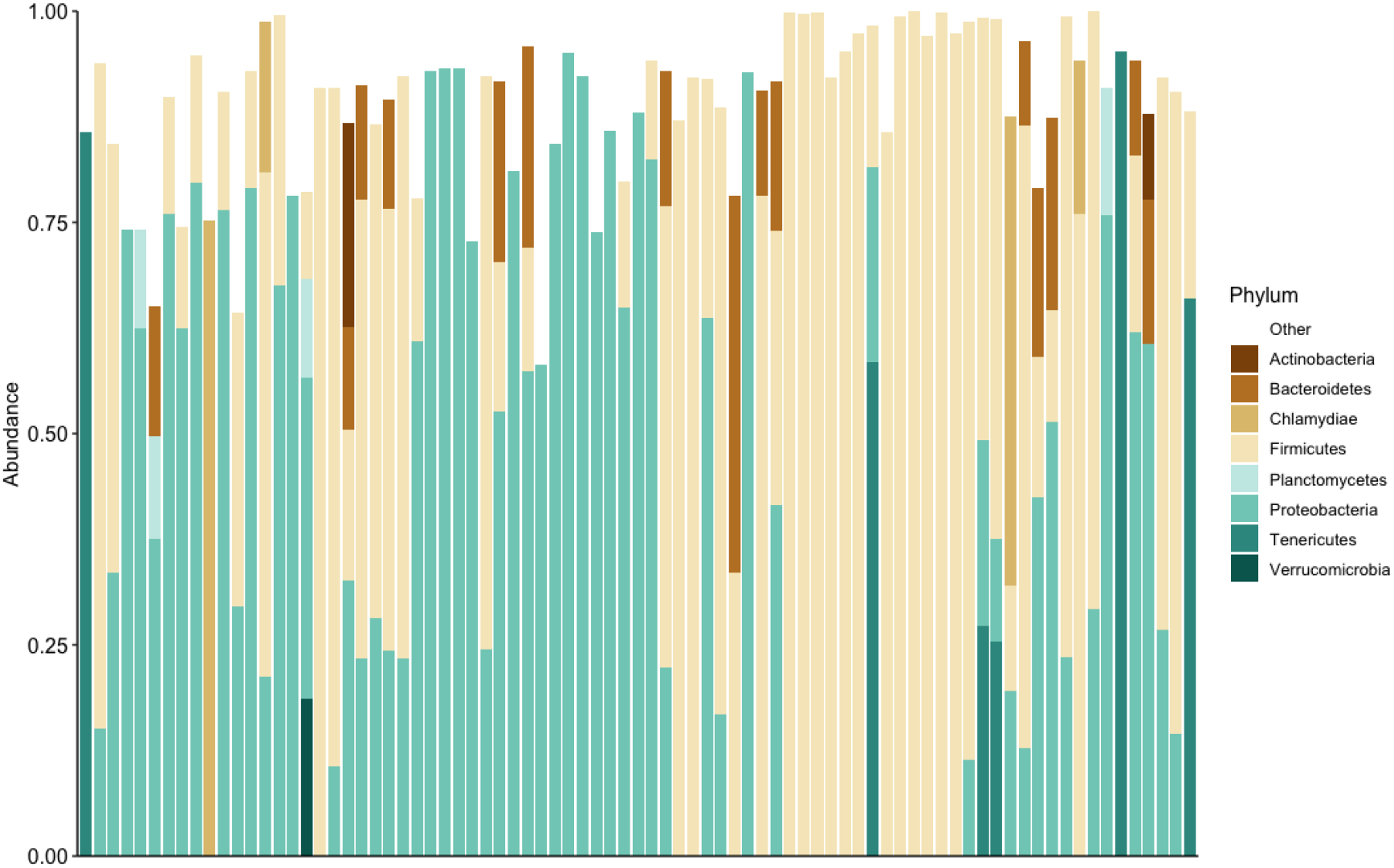
Relative abundance of the 8 most prominent phyla (>=10% relative abundance). Each vertical bar corresponds to a nestling at Day 8. “Other” group represents all low abundance phyla as white space.

### Nestling weight

The time-lagged analysis indicated that weight at D15 was negatively related to alpha diversity at D8 (Figure 2a, Table 1, Table 4), controlling for weight at D8 (Table 1). Neither habitat, lay date or brood size had detectable influence in this analysis (Table 1). Cross-sectional analyses indicated no relationship between contemporary weight and diversity at either D8 or D15 (Figure 2b and 2c, Table 4, Table S1). Brood size was negatively related to weight at D8. There was no evidence for an effect of habitat on weight in D15 birds. The relative abundances of Proteobacteria and Firmicutes did not predict weight in any of these analyses.

**Figure 2.**
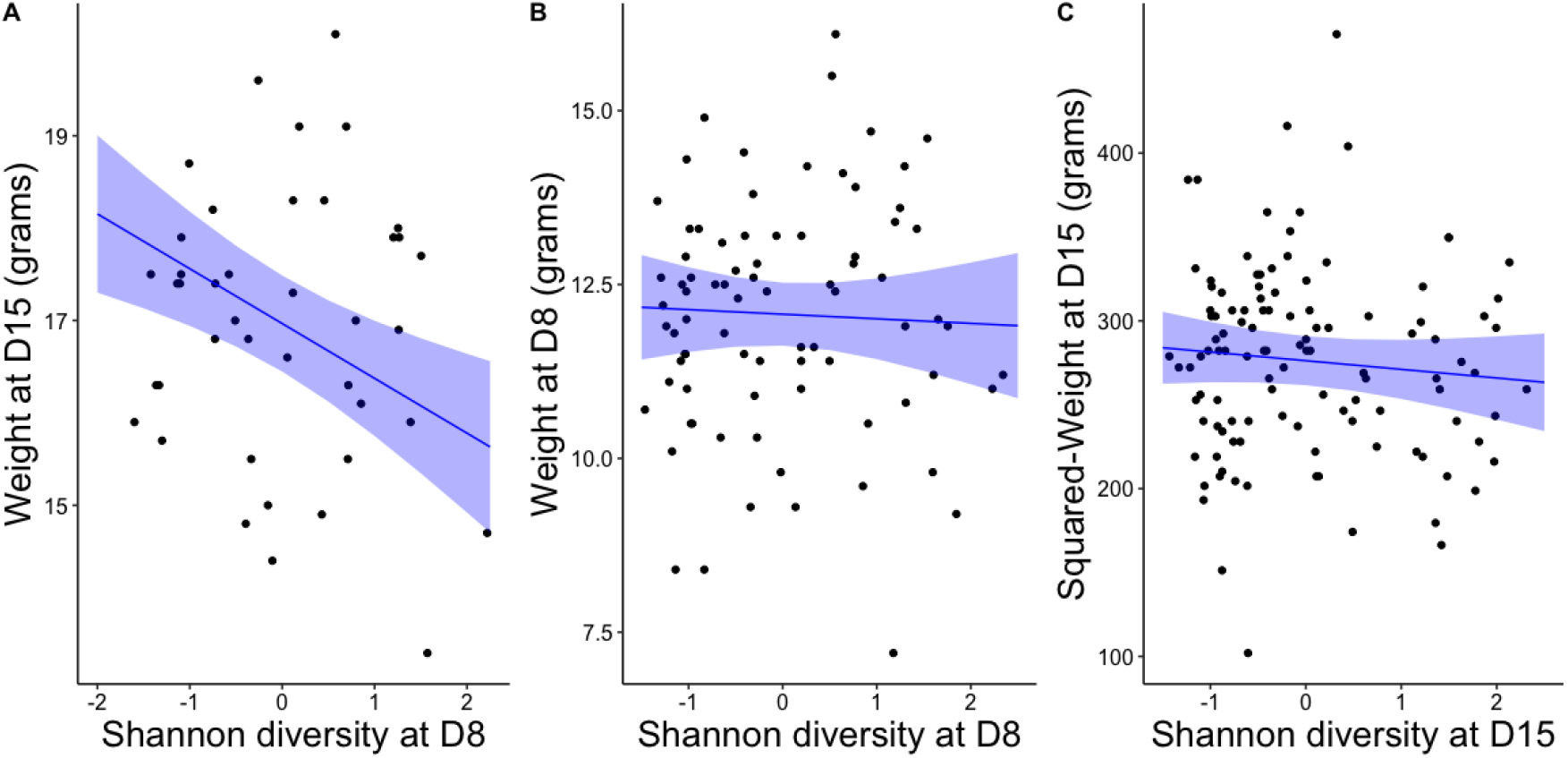
The effect of Shannon diversity on (a) time-lagged weight (i.e. Day 15 (D15)), (b) contemporary weight Day 8 (D8) and (c) contemporary weight D15. Black dots are individual data points, blue line is regression line with 95% CI (shaded blue). Shannon diversity is centred and scaled.

**Table 1.** GLMM output from time-lagged (Day-15) weight analyses. The effect of alpha diversity is reported for (a) Shannon diversity and (b) Chao1 diversity. * p < 0.05.

| Dependent~Independent variable | Estimate | Std. Error | df | Test statistic | P_estimate |
| --- | --- | --- | --- | --- | --- |
| **(a) Nestling weight (time-lagged)** |  |  |  |  |  |
| (Intercept) | 16.798 | 0.525 | 21.052 | 31.977 | <0.001 * |
| Shannon (Day-8) | -0.593 | 0.167 | 23.749 | -3.544 | 0.002 * |
| Habitat (deciduous) | 0.247 | 0.630 | 22.081 | 0.392 | 0.699 |
| Weight (Day-8) | 0.700 | 0.173 | 24.672 | 4.035 | <0.001 * |
| Lay Date (first egg) | 0.252 | 0.296 | 21.226 | 0.851 | 0.404 |
| Brood size (Day-8) | 0.168 | 0.273 | 21.991 | 0.616 | 0.544 |
| **(b) Nestling weight (time-lagged)** |  |  |  |  |  |
| (Intercept) | 16.961 | 0.475 | 20.856 | 35.681 | <0.001 * |
| Chao1 (Day-8) | -0.525 | 0.184 | 34.314 | -2.855 | 0.007 * |
| Habitat (deciduous) | 0.011 | 0.572 | 21.903 | 0.019 | 0.985 |
| Weight (Day-8) | 0.528 | 0.192 | 32.990 | 2.756 | 0.009 * |
| Lay Date (first egg) | 0.257 | 0.270 | 21.399 | 0.949 | 0.353 |
| Brood size (Day-8) | 0.121 | 0.250 | 22.841 | 0.483 | 0.634 |

Correlation tests indicate both Shannon (cor=0.33, t=2.17, df=39, p= 0.036) and Chao1 (cor=0.36, t=2.38, df=39, p=0.022) diversity measures were positively correlated within pairs.

### Nestling Survival

Survival was significantly related to weight at D8; the hazard ratio of 0.67 indicated that for every unit increase in weight, mortality risk decreased by a factor of 0.67. Survival was not influenced by lay date or brood size. Nestling survival to fledging was not predicted by Shannon’s diversity or Chao1 index at D8 (Table 2, Table 4). The relative abundance of Proteobacteria and Firmicutes at D8 also did not predict survival to fledging.

There was no clear evidence for the violation of the assumption of homogeneity of dispersion for survived/not-survived groups for Jaccard distance (Sum Squares= 0.009, F=3.295, p=0.081) or Bray-Curtis (Sum squares=0.006, F=1.973, p=0.164). Beta diversity did not differ between surviving and non-surviving nestlings for Jaccard (Sum Squares=0.563, F=1.233, p=0.172) or Bray-Curtis (Sum Squares=0.523, F=1.193, p=0.228), Table 4, and see Supporting Information Figure S2, Table S3.

**Table 2.** Determinants of mortality risk from Cox proportional hazards model. Positive coefficients indicate increased risk of mortality. HR = Hazard ratio, which is the exponentiated coefficients, and gives the ratio of the total number of observed to expected events. Note: all numeric covariates are centered and scaled. * p < 0.05.

| Dependent~Independent variable | Coefficient | HR | Standard<br>error | Test statistic (z) | P estimate |
| --- | --- | --- | --- | --- | --- |
| **Nestling mortality risk** |  |  |  |  |  |
| Shannon D8 | 0.021 | 1.021 | 0.327 | 0.07 | 0.95 |
| Weight D8 | -0.665 | 0.514 | 0.282 | -2.36 | 0.018 * |
| Lay Date (First egg) | 0.304 | 1.356 | 0.382 | 0.80 | 0.43 |
| Brood Size (Day-8) | 0.551 | 1.735 | 0.363 | 1.52 | 0.13 |
| **Nestling mortality risk** |  |  |  |  |  |
| Chao1 D8 | 0.314 | 1.368 | 0.283 | 1.11 | 0.27 |
| Weight D8 | -0.609 | 0.544 | 0.289 | -2.11 | 0.035 * |
| Lay Date (First egg) | 0.297 | 1.346 | 0.396 | 0.75 | 0.45 |
| Brood Size (Day-8) | 0.634 | 1.885 | 0.386 | 1.64 | 0.1 |

### Indicator species analysis

The indicator species with the highest indicator values associated with survival belonged to ASV46, ASV310 and ASV150 which all belong to the *Lactobacillaceae* family (Figure 3, Table 4, Table S4). The taxa with the highest indicator values associated with non-survival belonged to the ASV38, ASV1034, ASV895 which belong to *Enterobacteriaceae*, *Methylopilaceae* and *Acidothermaceae* families, respectively. The taxa indicative of non-survival and survival (p value <=0.05 and test statistic >=0.4) were plotted on a heatmap (Figure 3). Overall, specificity values were high (mean = 0.94±0.008 standard error) and fidelity values were low (0.21±0.008), suggesting that indicator species were typically only detected within a given group (i.e. survive/non-survive), but that not all birds within a group were recorded to have a given indicator species. All indicator species and test statistics (specificity values and indicator values) are reported in Supporting Information Table S4.

**Figure 3.**
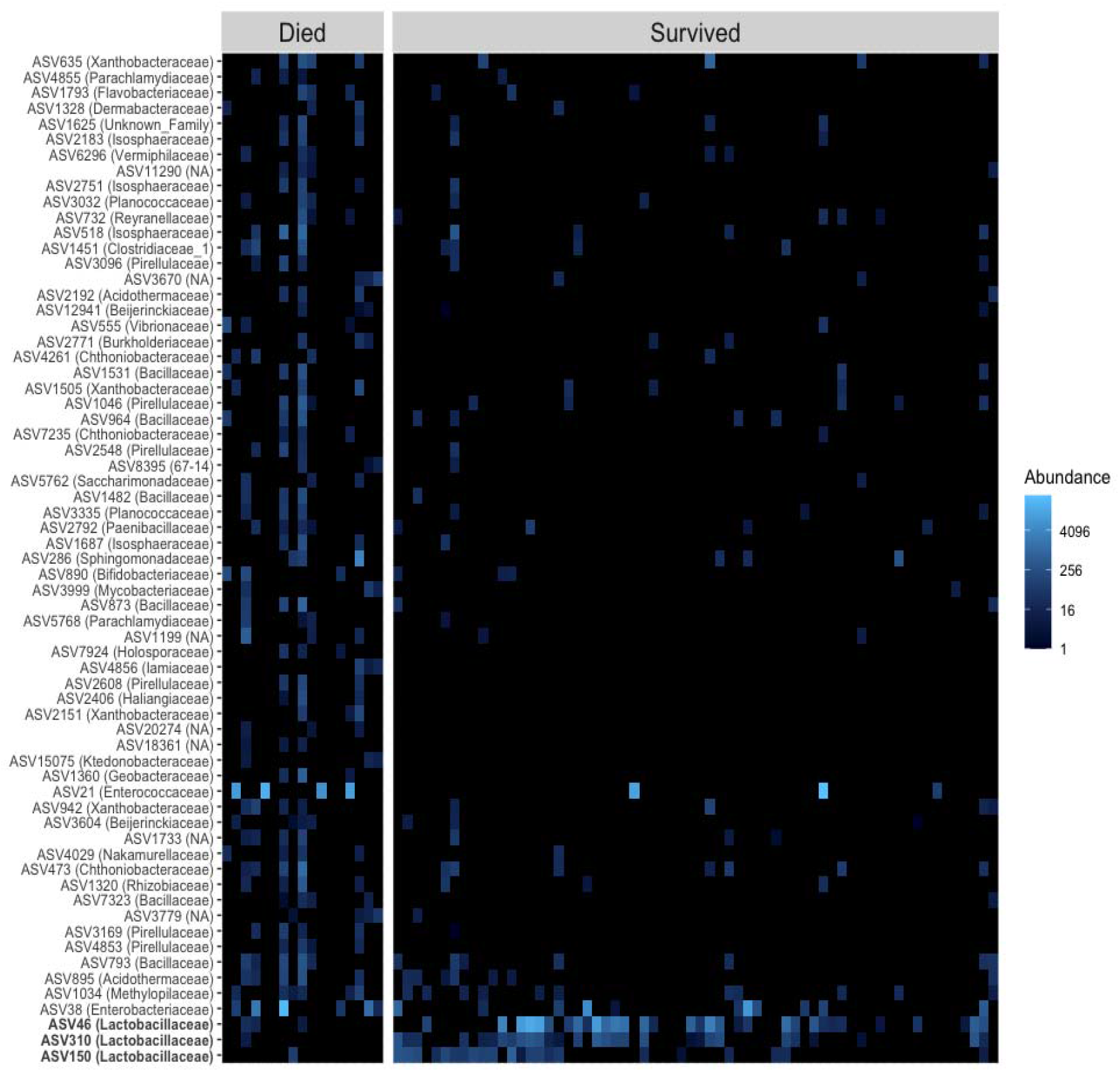
Heatmap of relative abundance of indicator species ASVs for survival. Individual birds are columns with separate panels for individuals that either died or survived. ASVs are in rows where taxa in brackets refer to the ASVs family-level taxonomic assignment, or NA if not resolved to family-level. Taxa associated with survival as opposed to non-survival are in bold. Lighter colours indicate greater abundance.

A posteriori analyses showed that a higher relative abundance of ASV46 (a *Lactobacillus* spp., and the taxa most strongly associated with survival) at D8 predicted a higher weight at D15. Relative abundance of ASV38 (a Rahnella sp., and the taxa most strongly associated with non-survival) was not associated with weight at D15 (Table 3, Table 4).

**Table 3.** A posteriori GLMM output from time-lagged weight analyses. The effect of indicator species on weight at D15 are reported for a) *Lactobacillus* sp. (ASV46) and b) Rahnella sp. (ASV38). * p <0.05

| Dependent/Independent variable | Estimate | Std.<br>Error | df | Test<br>statistic | P_estimate |
| --- | --- | --- | --- | --- | --- |
| **(a) D15 Weight (time-lagged)** |  |  |  |  |  |
| (Intercept) | 17.673 | 0.739 | 4.679 | 23.906 | <0.001 * |
| Abundance ***Lactobacillus* sp.**<br>**(ASV46)** | 1.042 | 0.367 | 8.601 | 2.839 | 0.02 * |
| Habitat (deciduous) | -1.946 | 1.007 | 5.126 | -1.933 | 0.11 |
| Weight (Day-8) | 0.835 | 0.340 | 11.184 | 2.460 | 0.031 * |
| Brood Size (Day-15) | -0.459 | 0.352 | 11.573 | -1.303 | 0.218 |
| Lay Date (First egg) | -0.457 | 0.389 | 7.245 | -1.175 | 0.277 |
| **(b) D15 Weight (time-lagged)** |  |  |  |  |  |
| (Intercept) | 16.473 | 0.656 | 2.000 | 25.111 | 0.002 * |
| Abundance ***Rahnella* sp.**<br>**(ASV38)** | 0.040 | 0.388 | 2.000 | 0.103 | 0.927 |
| Habitat (deciduous) | 2.908 | 1.320 | 2.000 | 2.203 | 0.158 |
| Weight (Day-8) | 5.398 | 1.579 | 2.000 | 3.419 | 0.076 |
| Brood Size (Day-15) | 1.690 | 0.660 | 2.000 | 2.558 | 0.125 |
| Lay Date (First Egg) | -1.069 | 1.113 | 2.000 | -0.960 | 0.438 |

**Table 4.** Summary of the relationship between microbiota and host performance.

| Microbiome metric | Survival | Non-survival | Weight (time-lagged) | Weight (contemporary) |
| --- | --- | --- | --- | --- |
| Alph diversity | NS | NS | (-) | NS |
| Beta diversity | NS | NS |  |  |
| Phylum-level abundance | NS | NS | NS | NS |
| Indicator species | (+) | (+) |  |  |
| *Lactobacillus* sp. (ASV 46) |  |  | (+) | NS |
| *Rahnella* sp. (ASV38) |  |  | NS | NS |

Significant relationships are shown as either positive (+), negative (-) or non significant (NS).

## Discussion

### Overall summary

We demonstrate that the gut microbiota correlates with nestling performance in a population of wild passerine birds. We found a negative association between weight at D15 and alpha diversity at D8. In other words, individuals with a low microbial diversity at D8 were heavier than expected at D15, given their weight at D8. Although alpha diversity at D8 was correlated with alpha diversity at D15, neither microbial alpha diversity nor phylum-level abundance predicted contemporary weight. Our exploratory analysis indicated that eight-day old nestlings with a greater relative abundance of specific *Lactobacillus* taxa were more likely to fledge, and our a posteriori analysis showed that for at least one of these taxa, greater relative abundance was specifically associated with increased host weight (relative to weight at D8). Notably, although we present potential links between gut microbiota and survival, and a proxy for survival (i.e. weight), the direction of these relationships or the microbial metrics that predict such relationships are variable within our own study (Table 4) and contrast with some, but not other studies. A lack of consensus on what aspects of microbial taxonomic composition predict host traits is prominent in the microbiome literature (Zaneveld et al., 2017). This may be due to broad statistical approaches that are correlative in nature, and therefore require manipulative studies to confirm relationships found here. Nevertheless, we discuss below how methodological approaches, critical developmental windows, environmental matching and the Anna Karenina Hypothesis may explain such differential findings across studies. Moreover, we suggest hypotheses worthy of future confirmatory testing to understand the effects of gut microbiota on the development of host phenotypes.

We show a negative relationship between alpha diversity and relative weight gain, over one nesting season. Experimentally reducing diversity of the microbiome through antibiotic administration has been shown to improve host growth (Banerjee et al., 2018; Kohl et al., 2018; Lan et al., 2005; Potti et al., 2002), possibly because antibiotics reduce bacterial load, diverting resources towards growth, as opposed to immune function, in a classic life-history trade-off (Rauw, 2012). Our study demonstrates that this negative relationship is present in the absence of microbiome manipulation, although we can only speculate as to the mechanisms responsible for limiting relative weight gain in nestlings with high gut microbial diversity. For example, a conventional gut microbiome compared to a germ-free one lowers absorption capacity of glucose and B-vitamins by increasing the thickness of the gut epithelium (Ford & Coates, 1971), and non-mutualistic microbes may create greater competition for nutrients (Wasielewski et al., 2016). While the diversity of the microbial community may be important for regulating the host’s future weight, some taxa may have disproportionate effects. At least one ASV, *Lactobacillus* sp., with the strongest potential association with survival, showed a positive, time-lagged association with host weight (relative to weight at D8). Several *Lactobacillus* spp. have beneficial, probiotic effects on the host (Angelakis & Raoult, 2010; Banerjee et al., 2018; Gomes et al., 2012; Vásquez et al., 2012), though this is certainly not the case for all *Lactobacillus* spp. Nevertheless, we can speculate as to why Lactobacillus may confer benefits to great tit nestlings, and encourage confirmatory research to test such hypotheses. For example, some strains of Lactobacillus sakei (ASV150) are known to produce antimicrobial peptides which inhibit a known pathogen (Gomes et al., 2012). Therefore we hypothesise that the presence of *Lactobacillus sakei*, which was associated with survival, may serve as an anti-pathogen in great tit nestlings. More generally, *Lactobacillus* spp. produce metabolic byproducts such as short chain fatty acids that can improve host energy metabolism (LeBlanc et al., 2017) and modulate immune response (Ratajczak et al., 2019). For example, the SCFA butyrate acts as an anti-inflammation agent, enhances intestinal barrier function and modulates the production of mucosal immune products in mammals (see review: Liu et al. (2018)). Future studies should look to shotgun sequencing to identify specific bacterial strains and genes for functional profiling and for developing probiotic treatments. This could be followed up by manipulative studies to identify the causal relationships between ASV and host traits, such as isolating and culturing ASV46 to test its potential probiotic effects when administered to nestlings (Davidson et al., 2020; Gatesoupe, 1999; Holzapfel & Schillinger, 2002).

Although a growing number of studies have demonstrated that the microbiome is an important predictor of host weight, the microbial profiles and the direction of this relationship vary across species and studies (Kohl et al., 2018; Potti et al., 2002; Teyssier, Lens, et al., 2018; Videvall et al., 2019). For example, a negative relationship between alpha diversity and future weight gain was found in juvenile ostriches (Videvall et al., 2019), whereas a positive relationship between alpha diversity and contemporary body condition was reported in a different population of great tit nestlings (Teyssier, Lens, et al., 2018). The contrasting results reported in these studies and our own may be due to variations in the microbiome sampling method and storage solutions (i.e. cloacal versus faecal, see Videvall et al. (2017); storing solutions see Vargas-Pellicer et al. (2019)). Moreover, our study and that of others are limited to a single nesting season, and any general patterns of the gut microbiota on host traits may differ across years according to local conditions such as food availability (see below for discussion of Predictive Adaptive Response hypothesis). Therefore it is likely that differences in local environments (Gillingham et al., 2019; Teyssier, Rouffaer, et al., 2018; Youngblut et al., 2019) and the temporal resolution of microbial and host traits dictate when and how microbiota impacts host phenotypes in birds (Videvall et al., 2019).

Time-lagged associations between the gut microbiota and host traits are expected to be especially important during developmental periods when large shifts in the establishing microbial community coincide with the programming of host biology (Cox et al., 2014; Hansen et al., 2012; Sudo et al., 2004), and in systems that are likely to experience abrupt changes in environmental inputs associated with gut microbiome, such as diet. The latter may be especially relevant in birds as their microbiomes are tightly linked with environmental variation, in contrast to mammals where the microbiome is primarily linked to phylogeny (Song et al., 2020). In ostriches, microbiome diversity predicted future weight gain, but the direction of this relationship switched after week 1 post-hatching, a time when ostriches no longer rely on an internal yolk as sustenance (Videvall et al., 2019). In the current study, alpha diversity at D8 predicted future weight at D15, whereas cross-sectional analyses were not sufficient to detect such a relationship. This time-lagged association may be owing to a critical developmental period in which the microbiome impacts on future phenotypes, a phenomenon that is prominent in clinical microbiome literature (Cox et al., 2014; Hansen et al., 2012; Sudo et al., 2004). The gut microbiome may also act as a mechanism for adaptive phenotypic change through its sensitivity to environmental variability, and its connection to host biological processes (Alberdi et al., 2016). Such a mechanism would also be in line with the ‘Predictive Adaptive Response hypothesis’, which posits that conditions during development cause phenotypic changes that serve to match individuals to their environments (Bateson et al., 2014; Crino & Breuner, 2015; Weber et al., 2018). For example, unpredictable food supply during the development stage can influence adult physiological and behavioral phenotypes (Spencer et al., 2009; Zimmer et al., 2013). Therefore, microbiome-mediated developmental plasticity may be adaptive in species where offspring benefit from a gut microbiome that primes the host for dietary shifts. Dietary shifts are common in tits where the relative frequency of insect taxa can change throughout the provisioning period (Arnold et al., 2007; García-Navas et al., 2012). Although Belgian great tit nestlings had the greatest improvement in body condition if they maintained a stable microbial diversity throughout development (Teyssier, Lens, et al., 2018), perhaps this may be because the environmental conditions at the time were also stable. In our study, it is difficult to differentiate between whether the gut microbiome is responding to dietary shifts as opposed to priming the host. Pinpointing if and when early life bacterial diversity programs host digestive and/or metabolic efficiency in wild animals, and the extent to which environmental variation, such as food availability and diet, impacts the developing microbiome will progress our understanding of microbial-mediated developmental mechanisms.

The Anna Karenina Hypothesis states that changes in an unhealthy or so-called “dysbiotic” microbiome can be stochastic rather than deterministic, making them difficult to detect statistically (Zaneveld et al., 2017). In other words, unhealthy individuals can have greater variation in their microbial community composition than healthy individuals, and rather than displaying a specific ‘unhealthy’ state, the microbiomes of diseased or stressed individuals can all look very different due to a breakdown in regulation. Although our study did not examine diseased individuals, as the original Anne Karenina Hypothesis was intended, this phenomenon may nevertheless explain why nestling survival was not associated with either alpha or beta diversity. In contrast to the Anne Karenina Hypothesis and our own study, disease-causing mortality in juvenile ostriches was associated with low alpha diversity, and beta diversity differed between diseased and non-diseased birds (Videvall et al., 2020). In the present study, although several indicator species were identified to be associated with birds that did or did not survive, many of these had low prevalence across nestlings. This is reflected in the high specificity values and low fidelity values, suggesting that although the presence of an indicator species in a nestling’s gut microbiome reflects a high probability of survival or non-survival, the same indicator species is not present across all surviving/non-surviving nestlings in our sample population. This distinction is important in, for example, conservation practices whereby microbes are used as markers for population viability (Trevelline et al., 2019; West et al., 2019). Our analysis suggests that in the case of non-survival, there are no microbes that are universally indicative of non-survival across birds, but rather the presence of one of several candidate microbes predicts a higher probability of mortality, which is consistent with the Anna Karenina Hypothesis. Alternatively, the gut microbiome does not predict survival in great tit nestlings at all, and the indicator species flagged in our study are an artifact of multiple testing without a correction due to the exploratory nature of this analysis. Therefore, whether these same taxa affect survival in other populations and/or other species of birds should be examined in independent confirmatory studies. For example, we hypothesise that ASVs reported for non-survival may be pathogenic to great tits, although we found no evidence in the literature to support this. Nevertheless, ASV1320, ASV2151, ASV3604 and ASV942 are members of the Rhizobiales order which contains genera that are well known human pathogens, and ASV5768 and ASV4855 are members of the Chlamydiales order which also contain pathogenic genera, though these genera were not detected in our study.

We also encourage further investigations into whether indicator taxa are the cause or consequence of host growth, or indeed any trait that may be associated with survival, such as glucocorticoids (Weber et al., 2018) and immune function (Hõrak et al., 1999). Manipulation of the gut microbiome alongside metabolomics or metagenomics will be the next step in providing insight into the causal direction and functional mechanisms of microbial variation and its relationship with host fitness. The latter is particularly important for identifying whether there is functional overlap between microbial taxa and the mechanisms by which these taxa affect host traits (Heintz-Buschart & Wilmes, 2018). Given the temporal importance of host-microbial relationships identified in this study and elsewhere, we anticipate the maturing microbiome will have important consequences on long-term fitness effects. More generally, our results highlight the importance of the gut microbiome in evolutionary processes as it has the potential to mediate rapid adaptation to environmental change and may also support conservation efforts as an indicator of individual and population health.

## Acknowledgements

Thanks to Jodie Crane and Iván de la Hera, for assistance with fieldwork. Paul Cotter, Fiona Crispie and Laura Finnegan from the Teagasc Sequencing facility for their role in relation to the 16S rRNA sequencing. For advice on bioinformatics, we thank Fiona Fouhy, Orla O’Sullivan and Calum Walsh. We are grateful to James Nichols, Ally Phillimore and Sarah Knowles for helpful discussions on molecular methodology, and to Enrico Pirotta for advice on statistical analyses. Thank you to Professor Jean-Michel Gaillard, the Associate Editor, and two anonymous referees for their constructive comments.

## Data accessibility

Our metadata is deposited at Dryad: doi:10.5061/dryad.bk3j9kd9g, and sequence data will be uploaded to a repository such as NCBI Sequence Read Archive, if the manuscript is published.

## Research and Animal Ethics

This study was conducted under licences from the Health Products Regulatory Authority (AE19130_P017), The National Parks and Wildlife Services (010/2016) and the British Trust for Ornithology, and permission from Coillte Forestry and private landowners. The research project received ethical approval from the Animal Welfare Body at University College Cork, and was in accordance with the ASAB (Association for the Study of Animal Behaviour) Guidelines for the Treatment of Animals in Behavioural Research and Teaching.

## Author contributions

GLD, CS, RPR and JLQ conceived the ideas and designed the methodology. GLD and MSR collected the field data. GLD, NW and CNJ carried out the DNA extraction and library prep. SES analysed the data. GLD and SES wrote the manuscript with input from all authors. All authors gave final approval for publication.

