## supplementary material for "A time-lagged association between the gut microbiome, nestling weight and nestling survival in wild great tits"

*Joint first authors

a. School of Biological, Earth and Environmental Sciences, Distillery Fields, North Mall, University College Cork, Cork, Ireland.
b. Department of Psychology, Downing Street, University of Cambridge, Cambridge, UK.
c. APC Microbiome Ireland, University College Cork, Cork, Ireland.
d. Teagasc Food Research Centre, Moorepark, Fermoy, Ireland.
e. Department of Integrative Biology, Oklahoma State University, USA

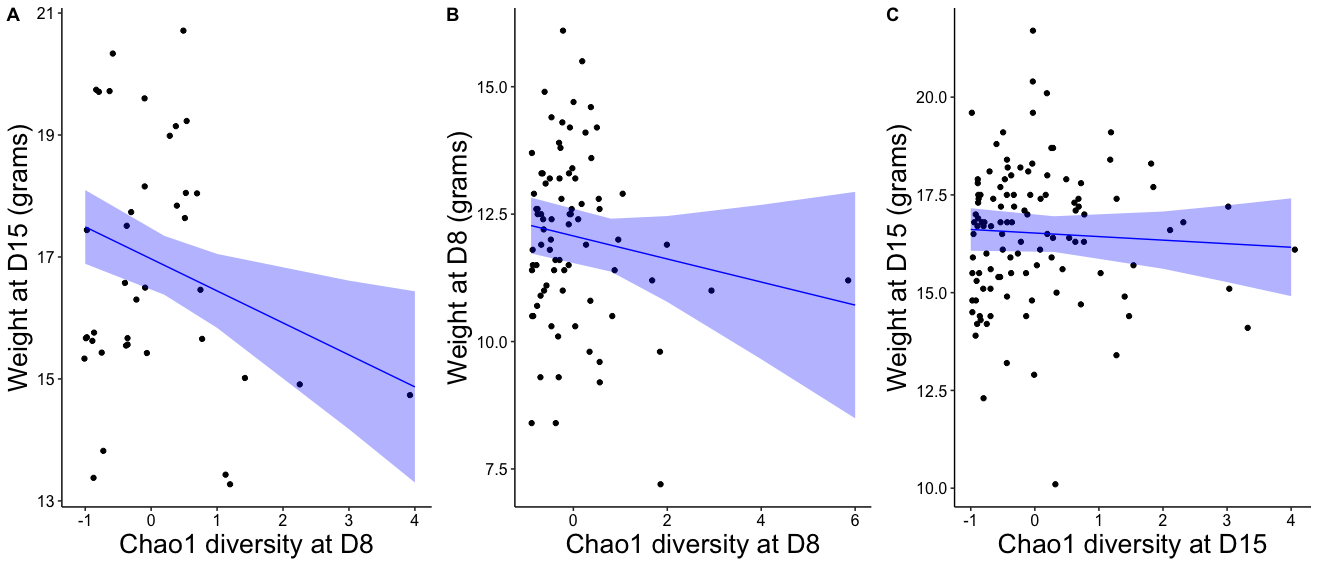

Figure S1. The effect of Chao1 diversity on (a) time-lagged weight (i.e. Day 15 (D15)), (b) contemporary weight D8 and (c) contemporary weight D15. Black dots are individual data points, blue line is regression line with 95% CI (shaded blue). Chao1 diversity is centred and scaled.

Table S1. GLMM outputs from contemporary weight analyses for (a) D8 and (b) D15. Alpha diversity is reported for (i) Shannon, and (ii) Chao1.* D15 weight was squared in order to allow model convergence. * p < 0.05

| **Dependent/Independent variable** | **Estimate** | **Std. Error** | **df** | **Test statistic** | **P_estimate** |
| --- | --- | --- | --- | --- | --- |
| **(a,i) Day-8 weight (contemporary)** | | | | | |
| (Intercept) | 12.666 | 0.507 | 30.714 | 24.985 | <0.001 * |
| Shannon | -0.066 | 0.194 | 73.965 | -0.341 | 0.734 |
| Habitat | -0.737 | 0.570 | 30.528 | -1.293 | 0.206 |
| Lay date (of first egg) | -0.031 | 0.240 | 27.262 | -0.130 | 0.897 |
| Brood size (when sampled) | -0.465 | 0.234 | 31.274 | -1.982 | 0.056 |
| **(a, ii) Day-8 weight (contemporary)** | | | | | |
| (Intercept) | 12.823 | 0.501 | 32.018 | 25.610 | <0.001 * |
| Chao1 | 0.000 | 0.000 | 74.167 | -1.237 | 0.22 |
| Habitat | -0.680 | 0.548 | 29.877 | -1.240 | 0.225 |
| Lay date (of first egg) | -0.024 | 0.229 | 26.155 | -0.104 | 0.918 |
| Brood size (when sampled) | -0.478 | 0.225 | 30.174 | -2.125 | 0.042 * |
| **(b, i) Day-15 weight* (contemporary)** | | | | | |
| (Intercept) | 304.157 | 15.979 | 39.296 | 19.035 | <0.001 * |
| Shannon | -5.148 | 5.148 | 107.825 | -1.000 | 0.32 |
| Habitat | -36.543 | 18.708 | 39.183 | -1.953 | 0.058 |
| Lay date (of first egg) | 4.627 | 7.612 | 40.062 | 0.608 | 0.547 |
| Brood size (when sampled) | -5.932 | 7.492 | 46.477 | -0.792 | 0.433 |
| **(b, ii) Day-15 weight (contemporary)** | | | | | |
| (Intercept) | 17.490 | 0.525 | 47.490 | 33.315 | <0.001 * |
| Chao1 | 0.000 | 0.000 | 104.772 | -0.612 | 0.542 |
| Habitat | -1.146 | 0.576 | 39.283 | -1.989 | 0.054 |
| Lay date (of first egg) | 0.105 | 0.234 | 39.904 | 0.450 | 0.655 |
| Brood size (when sampled) | -0.181 | 0.230 | 46.436 | -0.785 | 0.437 |

Table S2. PERMANOVA results from nestling beta diversity analysis. Models effect on (a) Jaccard and (b) Bray-Curtis distances, of D8 weight and survival status. * p < 0.05

| **Dependent/Independent variable** | **Df** | **SumOfSqs** | **R2** | **F** | **P_estimate** |
| --- | --- | --- | --- | --- | --- |
| **(a) Jaccard** | | | | | |
| D8 Weight | 1 | 0.495 | 0.014 | 1.085 | 0.329 |
| Survive | 1 | 0.563 | 0.016 | 1.233 | 0.172 |
| **(b) Bray-Curtis** | | | | | |
| D8 Weight | 1 | 0.495 | 0.014 | 1.085 | 0.329 |
| Survive | 1 | 0.563 | 0.016 | 1.233 | 0.172 |

Table S3. Effect on weight of phyla-level abundance for top 2 most abundant phyla, Proteobacteria and Firmicutes. (a-b) time-lagged weight, (c-d) D8 contemporary weight and (e-f) D15 contemporary weight. * p < 0.05

| **Dependent/Independent variable** | **Estimate** | **Std. Error** | **df** | **Test statistic** | **P_estimate** |
| --- | --- | --- | --- | --- | --- |
| **(a) Nestling weight** (time-lagged) | | | | | |
| (Intercept) | 17.093 | 0.506 | 20.181 | 33.777 | <0.001 * |
| Proteobacteria Abundance | 0.033 | 0.178 | 23.541 | 0.185 | 0.855 |
| Habitat | -0.179 | 0.609 | 21.275 | -0.294 | 0.771 |
| Weight (D8) | 0.650 | 0.216 | 33.602 | 3.005 | 0.005 * |
| Lay Date First | 0.178 | 0.286 | 20.597 | 0.623 | 0.54 |
| Brood Size (D15) | 0.087 | 0.271 | 23.469 | 0.320 | 0.752 |
| **(b) Nestling weight** (time-lagged) | | | | | |
| (Intercept) | 17.075 | 0.520 | 19.833 | 32.855 | <0.001 * |
| Firmicutes Abundance | 0.327 | 0.194 | 26.286 | 1.690 | 0.103 |
| Habitat | -0.174 | 0.624 | 20.792 | -0.279 | 0.783 |
| Weight (D8) | 0.568 | 0.204 | 30.418 | 2.788 | 0.009 * |
| Lay date (of first egg) | 0.108 | 0.297 | 20.824 | 0.363 | 0.72 |
| Brood Size (D15) | 0.220 | 0.284 | 23.839 | 0.775 | 0.446 |
| **(c) Day-8 weight (contemporary)** | | | | | |
| (Intercept) | 17.491 | 0.540 | 42.498 | 32.390 | <0.001 * |
| Proteobacteria Abundance | -0.001 | 0.005 | 4220.445 | -0.105 | 0.916 |
| Habitat | -1.166 | 0.584 | 42.561 | -1.997 | 0.052 |
| Lay date (of first egg) | 0.023 | 0.057 | 42.621 | 0.404 | 0.688 |
| Brood Size (D8) | -0.031 | 0.037 | 43.477 | -0.843 | 0.404 |
| **(d) Day-8 weight (contemporary)** | | | | | |
| (Intercept) | 12.790 | 0.519 | 37.065 | 24.624 | <0.001 * |
| Firmicutes Abundance | 0.167 | 0.132 | 220.964 | 1.261 | 0.209 |
| Habitat | -0.709 | 0.568 | 32.462 | -1.248 | 0.221 |
| Lay date (of first egg) | -0.048 | 0.159 | 29.880 | -0.301 | 0.765 |
| Brood Size (D8) | -0.540 | 0.285 | 28.342 | -1.895 | 0.068 |
| **(e) Day-15 weight (contemporary)** | | | | | |
| (Intercept) | 17.491 | 0.540 | 42.498 | 32.390 | <0.001 * |
| Proteobacteria Abundance | -0.001 | 0.005 | 4220.445 | -0.105 | 0.916 |
| Habitat | -1.166 | 0.584 | 42.561 | -1.997 | 0.052 |
| Lay date (of first egg) | 0.023 | 0.057 | 42.621 | 0.404 | 0.688 |
| Brood Size (D8) | -0.031 | 0.037 | 43.477 | -0.843 | 0.404 |
| **(f) Day-15 weight (contemporary)** | | | | | |
| (Intercept) | 12.790 | 0.519 | 37.065 | 24.624 | <0.001 * |
| Firmicutes Abundance | 0.167 | 0.132 | 220.964 | 1.261 | 0.209 |
| Habitat | -0.709 | 0.568 | 32.462 | -1.248 | 0.221 |
| Lay date (of first egg) | -0.048 | 0.159 | 29.880 | -0.301 | 0.765 |
| Brood Size (D8) | -0.540 | 0.285 | 28.342 | -1.895 | 0.068 |

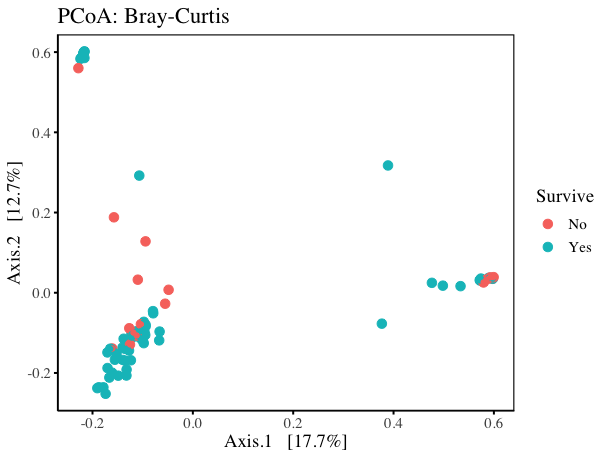

Figure S2. Principal Coordinate Analysis of Bray-Curtis distance matrix of Day-8 nestling samples, samples coloured by survival outcome.

Table S4. Indicator species output

| **ASV** | **Survival** | **Indicator Value** | **Specificity Value** | **Fidelity Value** | **p value** | **Phylum** | **Class** | **Order** | **Family** | **Genus** | **Species** |
| --- | --- | --- | --- | --- | --- | --- | --- | --- | --- | --- | --- |
| ASV1034 | N | 0.532 | 0.801 | 0.353 | 0.05 | Proteobacteria | Alphaproteobacteria | Rhizobiales | Methylopilaceae | Hansschlegelia | NA |
| ASV1046 | N | 0.41 | 0.952 | 0.176 | 0.05 | Planctomycetes | Planctomycetacia | Pirellulales | Pirellulaceae | Pir4_lineage | NA |
| ASV11290 | N | 0.407 | 0.938 | 0.176 | 0.015 | Actinobacteria | Acidimicrobiia | Microtrichales | NA | NA | NA |
| ASV1199 | N | 0.419 | 0.994 | 0.176 | 0.005 | Patescibacteria | Saccharimonadia | Saccharimonadales | NA | NA | NA |
| ASV12941 | N | 0.408 | 0.945 | 0.176 | 0.015 | Proteobacteria | Alphaproteobacteria | Rhizobiales | Beijerinckiaceae | Roseiarcus | NA |
| ASV1320 | N | 0.474 | 0.954 | 0.235 | 0.025 | Proteobacteria | Alphaproteobacteria | Rhizobiales | Rhizobiaceae | Mesorhizobium | NA |
| ASV1328 | N | 0.405 | 0.928 | 0.176 | 0.015 | Actinobacteria | Actinobacteria | Micrococcales | Dermabacteraceae | Brachybacterium | massiliense |
| ASV1360 | N | 0.42 | 1 | 0.176 | 0.005 | Proteobacteria | Deltaproteobacteria | Desulfuromonadales | Geobacteraceae | Geobacter | NA |
| ASV1451 | N | 0.408 | 0.943 | 0.176 | 0.04 | Firmicutes | Clostridia | Clostridiales | Clostridiaceae_1 | Clostridium_sensu_stricto_1 | disporicum |
| ASV1482 | N | 0.413 | 0.968 | 0.176 | 0.03 | Firmicutes | Bacilli | Bacillales | Bacillaceae | Bacillus | NA |
| ASV150 | Y | 0.549 | 0.877 | 0.344 | 0.05 | Firmicutes | Bacilli | Lactobacillales | Lactobacillaceae | Lactobacillus | sakei |
| ASV1505 | N | 0.41 | 0.952 | 0.176 | 0.03 | Proteobacteria | Alphaproteobacteria | Rhizobiales | Xanthobacteraceae | NA | NA |
| ASV15075 | N | 0.42 | 1 | 0.176 | 0.01 | Chloroflexi | Ktedonobacteria | Ktedonobacterales | Ktedonobacteraceae | NA | NA |
| ASV1531 | N | 0.409 | 0.949 | 0.176 | 0.015 | Firmicutes | Bacilli | Bacillales | Bacillaceae | Bacillus | niacini |
| ASV1625 | N | 0.405 | 0.928 | 0.176 | 0.025 | Proteobacteria | Gammaproteobacteria | Gammaproteobacteria_Incertae_Sedis | Unknown_Family | Acidibacter | NA |
| ASV1687 | N | 0.415 | 0.975 | 0.176 | 0.02 | Planctomycetes | Planctomycetacia | Isosphaerales | Isosphaeraceae | Singulisphaera | NA |
| ASV1733 | N | 0.463 | 0.912 | 0.235 | 0.035 | Proteobacteria | Alphaproteobacteria | Elsterales | NA | NA | NA |
| ASV1793 | N | 0.404 | 0.926 | 0.176 | 0.02 | Bacteroidetes | Bacteroidia | Flavobacteriales | Flavobacteriaceae | Myroides | NA |
| ASV18361 | N | 0.42 | 1 | 0.176 | 0.025 | Patescibacteria | Saccharimonadia | Saccharimonadales | NA | NA | NA |
| ASV20274 | N | 0.42 | 1 | 0.176 | 0.005 | Patescibacteria | Saccharimonadia | Saccharimonadales | NA | NA | NA |
| ASV21 | N | 0.423 | 0.759 | 0.235 | 0.03 | Firmicutes | Bacilli | Lactobacillales | Enterococcaceae | Catellicoccus | NA |
| ASV2151 | N | 0.42 | 1 | 0.176 | 0.02 | Proteobacteria | Alphaproteobacteria | Rhizobiales | Xanthobacteraceae | NA | NA |
| ASV2183 | N | 0.407 | 0.937 | 0.176 | 0.03 | Planctomycetes | Planctomycetacia | Isosphaerales | Isosphaeraceae | Singulisphaera | NA |
| ASV2192 | N | 0.408 | 0.945 | 0.176 | 0.015 | Actinobacteria | Actinobacteria | Frankiales | Acidothermaceae | Acidothermus | NA |
| ASV2406 | N | 0.42 | 1 | 0.176 | 0.015 | Proteobacteria | Deltaproteobacteria | Myxococcales | Haliangiaceae | Haliangium | NA |
| ASV2548 | N | 0.41 | 0.954 | 0.176 | 0.045 | Planctomycetes | Planctomycetacia | Pirellulales | Pirellulaceae | NA | NA |
| ASV2608 | N | 0.42 | 1 | 0.176 | 0.015 | Planctomycetes | Planctomycetacia | Pirellulales | Pirellulaceae | Pir4_lineage | NA |
| ASV2751 | N | 0.407 | 0.939 | 0.176 | 0.025 | Planctomycetes | Planctomycetacia | Isosphaerales | Isosphaeraceae | Aquisphaera | NA |
| ASV2771 | N | 0.409 | 0.947 | 0.176 | 0.03 | Proteobacteria | Gammaproteobacteria | Betaproteobacteriales | Burkholderiaceae | Xylophilus | NA |
| ASV2792 | N | 0.414 | 0.728 | 0.235 | 0.035 | Firmicutes | Bacilli | Bacillales | Paenibacillaceae | Paenibacillus | xylanilyticus |
| ASV286 | N | 0.415 | 0.976 | 0.176 | 0.03 | Proteobacteria | Alphaproteobacteria | Sphingomonadales | Sphingomonadaceae | Novosphingobium | NA |
| ASV3032 | N | 0.407 | 0.939 | 0.176 | 0.025 | Firmicutes | Bacilli | Bacillales | Planococcaceae | Sporosarcina | NA |
| ASV3096 | N | 0.408 | 0.943 | 0.176 | 0.05 | Planctomycetes | Planctomycetacia | Pirellulales | Pirellulaceae | NA | NA |
| ASV310 | Y | 0.67 | 0.992 | 0.453 | 0.01 | Firmicutes | Bacilli | Lactobacillales | Lactobacillaceae | Lactobacillus | NA |
| ASV3169 | N | 0.485 | 0.998 | 0.235 | 0.01 | Planctomycetes | Planctomycetacia | Pirellulales | Pirellulaceae | NA | NA |
| ASV3335 | N | 0.414 | 0.971 | 0.176 | 0.025 | Firmicutes | Bacilli | Bacillales | Planococcaceae | Chungangia | NA |
| ASV3604 | N | 0.441 | 0.825 | 0.235 | 0.04 | Proteobacteria | Alphaproteobacteria | Rhizobiales | Beijerinckiaceae | Methylobacterium | NA |
| ASV3670 | N | 0.408 | 0.944 | 0.176 | 0.01 | Cyanobacteria | Oxyphotobacteria | NA | NA | NA | NA |
| ASV3779 | N | 0.478 | 0.969 | 0.235 | 0.005 | NA | NA | NA | NA | NA | NA |
| ASV38 | N | 0.565 | 0.904 | 0.353 | 0.045 | Proteobacteria | Gammaproteobacteria | Enterobacteriales | Enterobacteriaceae | Rahnella | NA |
| ASV3999 | N | 0.416 | 0.981 | 0.176 | 0.01 | Actinobacteria | Actinobacteria | Corynebacteriales | Mycobacteriaceae | Mycobacterium | NA |
| ASV4029 | N | 0.467 | 0.927 | 0.235 | 0.01 | Actinobacteria | Actinobacteria | Frankiales | Nakamurellaceae | Nakamurella | NA |
| ASV4261 | N | 0.409 | 0.949 | 0.176 | 0.01 | Verrucomicrobia | Verrucomicrobiae | Chthoniobacterales | Chthoniobacteraceae | Candidatus_Udaeobacter | NA |
| ASV46 | Y | 0.706 | 0.997 | 0.5 | 0.025 | Firmicutes | Bacilli | Lactobacillales | Lactobacillaceae | Lactobacillus | NA |
| ASV473 | N | 0.468 | 0.932 | 0.235 | 0.05 | Verrucomicrobia | Verrucomicrobiae | Chthoniobacterales | Chthoniobacteraceae | Candidatus_Udaeobacter | NA |
| ASV4853 | N | 0.485 | 1 | 0.235 | 0.005 | Planctomycetes | Planctomycetacia | Pirellulales | Pirellulaceae | NA | NA |
| ASV4855 | N | 0.404 | 0.923 | 0.176 | 0.05 | Chlamydiae | Chlamydiae | Chlamydiales | Parachlamydiaceae | Neochlamydia | NA |
| ASV4856 | N | 0.42 | 1 | 0.176 | 0.01 | Actinobacteria | Acidimicrobiia | Microtrichales | Iamiaceae | Iamia | NA |
| ASV518 | N | 0.408 | 0.941 | 0.176 | 0.05 | Planctomycetes | Planctomycetacia | Isosphaerales | Isosphaeraceae | Aquisphaera | NA |
| ASV555 | N | 0.408 | 0.946 | 0.176 | 0.04 | Proteobacteria | Gammaproteobacteria | Vibrionales | Vibrionaceae | Vibrio | NA |
| ASV5762 | N | 0.412 | 0.962 | 0.176 | 0.005 | Patescibacteria | Saccharimonadia | Saccharimonadales | Saccharimonadaceae | NA | NA |
| ASV5768 | N | 0.418 | 0.989 | 0.176 | 0.02 | Chlamydiae | Chlamydiae | Chlamydiales | Parachlamydiaceae | Neochlamydia | NA |
| ASV6296 | N | 0.407 | 0.938 | 0.176 | 0.02 | Dependentiae | Babeliae | Babeliales | Vermiphilaceae | NA | NA |
| ASV635 | N | 0.402 | 0.687 | 0.235 | 0.025 | Proteobacteria | Alphaproteobacteria | Rhizobiales | Xanthobacteraceae | Afipia | NA |
| ASV7235 | N | 0.41 | 0.954 | 0.176 | 0.005 | Verrucomicrobia | Verrucomicrobiae | Chthoniobacterales | Chthoniobacteraceae | Chthoniobacter | NA |
| ASV732 | N | 0.407 | 0.941 | 0.176 | 0.05 | Proteobacteria | Alphaproteobacteria | Reyranellales | Reyranellaceae | Reyranella | NA |
| ASV7323 | N | 0.477 | 0.966 | 0.235 | 0.01 | Firmicutes | Bacilli | Bacillales | Bacillaceae | Bacillus | NA |
| ASV7924 | N | 0.42 | 1 | 0.176 | 0.005 | Proteobacteria | Alphaproteobacteria | Holosporales | Holosporaceae | NA | NA |
| ASV793 | N | 0.514 | 0.897 | 0.294 | 0.025 | Firmicutes | Bacilli | Bacillales | Bacillaceae | Bacillus | NA |
| ASV8395 | N | 0.411 | 0.958 | 0.176 | 0.02 | Actinobacteria | Thermoleophilia | Solirubrobacterales | 67-14 | NA | NA |
| ASV873 | N | 0.416 | 0.983 | 0.176 | 0.025 | Firmicutes | Bacilli | Bacillales | Bacillaceae | Bacillus | NA |
| ASV890 | N | 0.415 | 0.976 | 0.176 | 0.015 | Actinobacteria | Actinobacteria | Bifidobacteriales | Bifidobacteriaceae | Bifidobacterium | bifidum |
| ASV895 | N | 0.522 | 0.926 | 0.294 | 0.035 | Actinobacteria | Actinobacteria | Frankiales | Acidothermaceae | Acidothermus | NA |
| ASV942 | N | 0.434 | 0.802 | 0.235 | 0.045 | Proteobacteria | Alphaproteobacteria | Rhizobiales | Xanthobacteraceae | NA | NA |
| ASV964 | N | 0.41 | 0.953 | 0.176 | 0.035 | Firmicutes | Bacilli | Bacillales | Bacillaceae | Bacillus | NA |

**Microbiome and fitness in great tits: Supplementary methods**

***DNA Extractions***

Prior to extraction, the samples were centrifuged at 13,000 rpm for 3 minutes, and the ethanol was removed using a pipette. Samples were then allowed to dry in a sterile fume hood for 2-3 hours, or until visibly dry. Total DNA was extracted from the dried faecal contents of all birds using the Qiagen QIAamp DNA Stool Kit, following the "Isolation of DNA from Stool for Pathogen Detection" protocol (June 2012 edition), with some modifications following Zeale et al. (2011) and custom modifications to accommodate dried avian faeces. Briefly, a 0.10 - 0.20 g aliquot of faecal sample was added to 1.4 ml of buffer ASL supplied in the kit together with 0.4g of zirconia beads. The samples were homogenised for 1 minute using the Biospec Minibeadbeater. The suspension was then heated for 30 minutes at 70˚C following the addition of Proteinase K. Following centrifugation, 1 InhibitEx tablet was added to 1.2 ml of the supernatant and the suspension was incubated for 1 minute at room temperature to allow inhibitors to absorb to the InhibitEx matrix. The samples were centrifuged twice and 20 µl of proteinase K was added to 400 µl of the supernatant along with 400 µl of AL. The suspension was homogenised and incubated at 70˚C for 15 minutes. 400 µl of ethanol (96-100\%) was added to the lysate and mixed well by vortexing. Subsequent removal of RNA, protein and purification of the DNA was completed according to the manufacturer's instructions. DNA was then stored at -20˚C.

***Illumina MiSeq sequencing***

The V3-V4 variable region of the 16S rRNA gene was amplified from the DNA extracts using the 16S metagenomic sequencing library protocol (Illumina). The DNA was amplified with primers specific to the V3-V4 region of the 16S rRNA gene which also incorporates the Illumina overhang adaptor (Forward primer 5' TCGTCGGCAGCGTCAGATGTGTATAAGA- GACAGCCTACGGGNGGCWGCAG; reverse primer 5' GTCTCGTGGGCTCGGAGATGTGTATAA- GAGACA GGACTACHVGGGTATCTAATCC). Each PCR reaction contained 23 µl DNA template, 1 µl forward primer (10 µM), 1 µl reverse primer (10 µM) and 25 µl 2X Kapa HiFi Hotstart ready mix (Roche, Ireland), to a final volume of 50 µl. PCR amplification was carried out as follows: heated lid 110°, 95°C x 3mins, 30 cycles of 95°C x 30s, 55°C x 30s, 72°C x 30s, then 72°C x 5mins and held at 4°C. PCR products were visualised using gel electrophoresis (1X TAE buffer, 1.5\% agarose, 100V). Successful PCR products were cleaned using AMPure XP magnetic bead based purification (Labplan, Dublin, Ireland). A second PCR reaction was completed on the purified DNA (5µl) to index each of the samples, allowing samples to be pooled for sequencing on the one flow cell and subsequently demultiplexed for analysis. Two indexing primers (Illumina Nextera XT indexing primers, Illumina, Sweden) were used per sample. Each PCR reaction contained 5µl index 1 primer (N7xx), 5µl index 2 primer (S5xx), 25µl 2x Kapa HiFi Hot Start Ready mix, 10µl PCR grade water. PCRs were completed as described above, but only 8 amplification cycles were completed instead of 30. PCR products were visualised using gel electrophoresis and subsequently cleaned (as described above). Samples were quantified using the Qubit (Bio-Sciences, Dublin, Ireland), along with the broad range DNA quantification assay kit (Bio-Sciences) and samples were then pooled in an equimolar fashion. The pooled sample was run on the Agilent Bioanalyser for quality analysis prior to sequencing. The sample pool was prepared following Illumina guidelines. Samples were sequenced on the MiSeq sequencing platform (Clinical Microbiomics, Denmark), using a 2 x 300 cycle kit, following standard Illumina sequencing protocols.
